# Sensory nutrition: The role of taste in the reviews of commercial food products

**DOI:** 10.1101/662585

**Authors:** Danielle. R. Reed, Joel D. Mainland, Charles J. Arayata

**Affiliations:** Monell Chemical Senses Center, Philadelphia, PA 19104

**Keywords:** taste, sweetness, bitterness, consumers, segmentation

## Abstract

Many factors play a role in choosing what to eat or drink. Here we explored the role of sensation to explain these differences, drawing on consumer reviews for commercially available food products sold through an online retailer. We analyzed 393,568 unique food product reviews from Amazon customers with a total of 256,043 reviewers rating 67,553 products. Taste-associated words were mentioned more than words associated with cost, food texture, customer service, nutrition, smell, or those referring to the trigeminal senses, e.g., spicy. We computed the overall number of reviews that mentioned taste qualities: the word *taste* was mentioned in over 30% of the reviews (N= 142,768), followed by *sweet* (10.7%, N=42,315), *bitter* (2.9%, N=11,424), *sour* (2.1%, N=8,252), and *salty* (1.4%, N=5,688). We identified 38 phrases used to describe the evaluation of sweetness, finding that ‘too sweet’ was used in nearly 0.8% of the reviews and oversweetness was mentioned over 25 times more often than under-sweetness. We then focused on ‘polarizing’ products, those that elicited a wide difference of opinion (as measured by the ranges of the product ratings). Using the products that had more than 50 reviews, we identified the top 10 most polarizing foods (i.e., those with the largest standard deviation in ratings) and provide representative comments about the polarized taste experience of consumers. Overall, these results support the primacy of taste in real-world food ratings and individualized taste experience, such as whether a product is ‘too sweet’. Analysis of consumer review data sets can provide information about purchasing decisions and customer sensory responses to particular commercially available products and represents a promising methodology for the emerging field of sensory nutrition.

---

’Sensory nutrition’ is a research area examining how sensation affects what an animal (person) chooses to eat or drink and how these sensory-motivated choices affect their nutritional health. However, many studies that fall into the ‘sensory-nutrition’ study arena analyze these processes under artificial circumstances, e.g., one meal or one snack in the laboratory. Here we capitalize on real human behavior in a large arena of food choice, an on-line retailer that offers thousands of choices, examining the importance of sensory experience in the written comments of those who have purchased foods and drinks.

Sensory nutrition as a research area has become more essential because of the increasing realization that human foods have become ‘hyperpalatable’, engineered to make them so desirable from a sensory perspective that they are hard to resist and the overeating of these foods leads to obesity and other disease associated with overconsumption. However, while taste is often studied in simplified foods systems, like sugar dissolved in plain water [1], real-world foods are rarely evaluated for taste in a choice context in laboratories studies, although these types of studies are commonly done to evaluate products for the marketplace. Most broadly, the assertion is that the taste of foods drives overconsumption but that people differ in their perception. The larger question is whether personal differences in taste experience for different foods drive overconsumption of that food, and can ultimately predict who will not only like a certain food, but perhaps who cannot resist that food and why.

We took a step toward this larger question by analyzing how the sense of taste factors into food ratings by examining reviews of commercial foods written by customers of an online retailer. These ratings contain both a text narrative about the food and a ‘star’ rating, from one to five, with five stars representing the highest score. We were interested in how often taste was mentioned by reviewers and which of the taste qualities were mentioned most often. We were also interested in the idea that certain products were more polarizing among reviewers, with extreme diversity—love it or hate it—in responses, reasoning that a list of polarizing products might be a tool for future research to understand food choice. We extract and present reviewer comments about polarizing food products to illustrate what role taste played in creating the diverse viewpoints. We contrasted taste-specific works with those for texture (e.g., lumpy, creamy, soft, hard) and odor (e.g., smell) as well as price and customer service to see what sensory words were more common among reviewers.

## Methods

### Data and its structure

We obtained the data set through the open-source data competition site Kaggle (www.kaggle.com), where the data are offered freely to all under a Creative Commons public license. We performed all analyses in R (version 3.5.2) [2] and made this R script available on Github (https://github.com/joelmainland/Taste_is_king). The data contained ten variables: *Id* (each review has a unique identifier), *ProductId* (unique identifier for the product), *UserId* (identifier for the user), *Profile Name* (the self-assigned user name), *HelpfulnessNumerator* (the number of people who found the review helpful), *HelpfulnessDenominator* (the number of people who found the review helpful or not), *Score* (rating of the product on a 1 to 5 scale, with 5 being best and 1 being worst), *Time* (time of day the review was submitted), *Summary* (brief review), and *Text* (full review). The reviews were submitted over a ten-year period ending in October 2012. There were 568,454 reviews but some were duplicated (same text for similar products) and were removed prior to analysis. Analysis of the de-duplicated data indicated there were a total of 393,568 reviews of 67,553 products by 256,043 unique reviewers (as defined by unique reviewer IDs; however, this does not ensure absolute uniqueness because the same person might have more than one ID. All reviewers were ‘verified purchasers*,’* meaning the online retailer (Amazon) had a record that the reviewer purchased the food item being reviewed.

### Word use analysis

We used the Word2Vec package to create a vector representation of words based on the distributional hypothesis, namely that words that appear in the same context share semantic meaning [3]. Likes and dislikes can arise because of food tastes, textures or odors, cost, nutrition, or quality of service. Thus, the vector representation was used to identify clusters of semantically similar words based on seed words from these categories (Supplementary Table 1). We counted the numbers of reviews that contained words from these clusters. In some cases we elected not to use individual odor descriptors because odors are often described using the word for the object producing the odor [4]., e.g., honey smells like honey, peaches smell like peaches.

Building on the results that we explain below, we also probed for more details about sweet (because it was the most commonly used taste word). We extracted all phrases using the word ‘sweet’ and cleaned the data by eliminating common but irrelevant uses of the word sweet, e.g., ‘sweet potato’. We next extracted 38 phrases that captured the majority of ways sweetness was discussed, and tallied the number of times the phrase was used, e.g., ‘too sweet’ versus ‘not too sweet’ and used to calculate the percent of the reviews which contained that phrase. The phrases were placed into one of three categories: oversweet, (e.g., cloying, sickeningly sweet), under-sweet (e.g., not sweet enough for me) and neutral. We then tabulated the percentage of comments within each of the three categories.

### Polarization

We extracted all food products that had 50 or more reviews and computed the standard deviation of the ratings (on the star scale). We refer to the foods with the largest standard deviation as ‘polarizing’ foods. We chose the top 10 polarizing foods for a more in-depth analysis. As we mentioned above, each product is identified by a unique ID number. To find out which foods were associated with which ID number, we automatically extracted data about the Amazon product page for each ID to obtain the product title. For each of the 10 most polarizing products, two readers evaluated the narrative portion of each review (*Summary* plus *Text*) and extracted representative comments about taste.

## Results

### Overview

We examined several global categories using seed words: taste and related words, price and related words, likewise customer service, texture, smell and trigeminal (e.g., spiciness). In this analysis, the predominant word used in reviews was ‘taste’ with over 30% of reviews using this word; fewer than those who mentioned ‘price’ (Table 1). To examine whether this method of generating words was valid, we compared this list of words obtained to those words used by sensory panels in the food industry to describe texture [5], finding substantial agreement, e.g., hard, rough. For smell, we elected not to use individual odor descriptors as seed words because odors are often named for the physical source [4] and we cannot easily differentiate when the word ‘coffee’ is used as a product description from when it is used to describe an odor. When the results are aggregated over the five categories, the results show that ‘taste’ is mentioned more often than texture, customer service, cost, health, smell, and trigeminal sensations (Figure 1). Reviewers mentioned *sweet* far more often than any other taste quality (10.75%), followed by *bitter* (2.90%), *savory* (0.27%), *sour* (2.10%), and *salty* (1.45%). *Umami* is a synonym for *savory* but a word that is rarely used by reviewers (0.02%).

**Table 1:** Word counts by category

| Word | Count | Percent | Category |
| --- | --- | --- | --- |
| taste | 121447 | 30.86 | Taste |
| price | 53327 | 13.55 | Price |
| order | 52279 | 13.28 | Customer Service |
| sweet | 42315 | 10.75 | Taste |
| light | 27329 | 6.94 | Texture |
| ship | 26640 | 6.77 | Customer Service |
| hard | 23253 | 5.91 | Texture |
| rough | 22058 | 5.60 | Texture |
| healthy | 19674 | 5.00 | Health |
| smell | 17457 | 4.44 | Smell |
| organic | 16239 | 4.13 | Health |
| texture | 15544 | 3.95 | Texture |
| diet | 13514 | 3.43 | Health |
| fine | 13475 | 3.42 | Texture |
| arrived | 12368 | 3.14 | Customer Service |
| deal | 12259 | 3.11 | Price |
| bitter | 11424 | 2.90 | Taste |
| cheaper | 9481 | 2.41 | Price |
| sour | 8252 | 2.10 | Taste |
| value | 7506 | 1.91 | Price |
| spicy | 6945 | 1.76 | Trigeminal |
| aroma | 6724 | 1.71 | Smell |
| salty | 5688 | 1.45 | Taste |
| service | 5613 | 1.43 | Customer Service |
| condition | 5264 | 1.34 | Customer Service |
| sale | 5053 | 1.28 | Price |
| healthier | 4375 | 1.11 | Health |
| safe | 3941 | 1.00 | Health |
| bland | 3889 | 0.99 | Trigeminal |
| kick | 3868 | 0.98 | Trigeminal |
| scent | 3232 | 0.82 | Smell |
| fruity | 2045 | 0.52 | Smell |
| odor | 1803 | 0.46 | Smell |
| tabasco | 572 | 0.15 | Trigeminal |
| spiciness | 395 | 0.10 | Trigeminal |

'Trigeminal' refers to the common chemical sense, e.g., burning, stinging, cooling.

We also probed in more detail about sweet and sweetness and identified 38 phrases used to indicate this property. We considered each phrase and its negation, e.g., ‘sweet enough’ versus ‘not sweet enough’ and parsed each phrase into one of three categories, over-sweet, under-sweet or neutral. The results were striking. Almost 1 percent of all reviews, regardless of food type, used the phrase ‘too sweet’ indicating that excessive sweetness is often mentioned. When evaluating the pattern of reviews that mention sweetness, over-sweetness was mentioned more than 25 times more often than under-sweetness (Table 2). See Supplemental Table 2 for a list of the 38 phrases, the counts, and percentage of time each phrase was used in all reviews.

**Table 2:** Sweet taste phrase use in reviews

| Count | Sweetness | Percent |
| --- | --- | --- |
| 7230 | Over | 56.2 |
| 5370 | Neutral | 41.7 |
| 268 | Under | 2.1 |

### Polarizing products

For this analysis, we excluded products with fewer than 50 reviews, reducing the number of reviews by roughly half (N=109,698) and the number of products from 67,533 to 908. We computed the standard deviation for each remaining product and ranked the products from highest to lowest standard deviation. After excluding products not intended for human consumption (e.g., pet food), we selected the top 10 products (Table 3). Standard deviations ranged from 1.82 (most polarizing) to 0.21 (least polarizing).

**Table 3:**
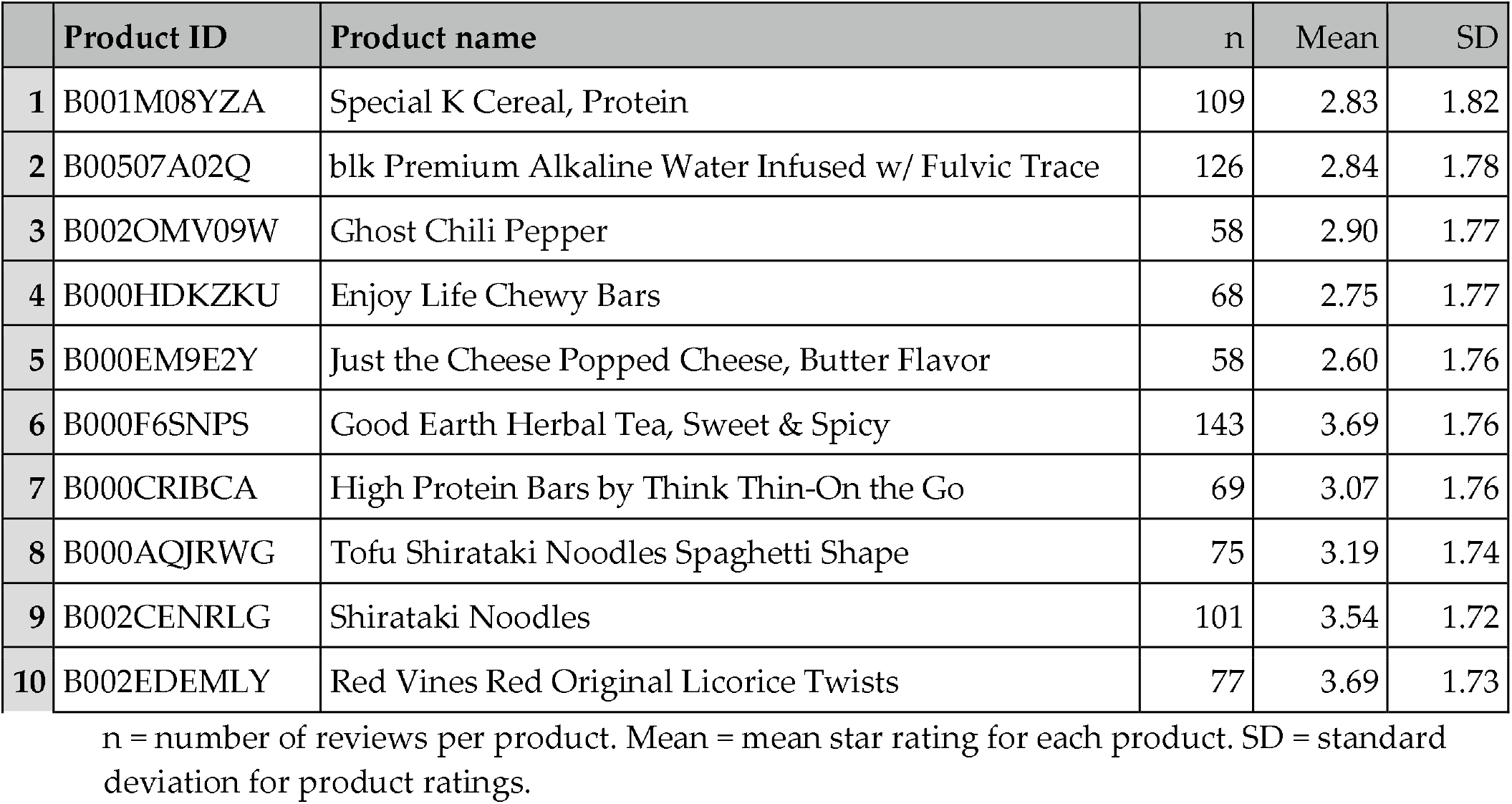
Products with the highest standard deviation in rating (polarization)

|  | Product ID | Product name | n | Mean | SD |
| --- | --- | --- | --- | --- | --- |
| 1 | B001M08YZA | Special K Cereal, Protein | 109 | 2.83 | 1.82 |
| 2 | B00507A02Q | blk Premium Alkaline Water Infused w/ Fulvic Trace | 126 | 2.84 | 1.78 |
| 3 | B002OMV09W | Ghost Chili Pepper | 58 | 2.90 | 1.77 |
| 4 | B000HDKZKU | Enjoy Life Chewy Bars | 68 | 2.75 | 1.77 |
| 5 | B000EM9E2Y | Just the Cheese Popped Cheese, Butter Flavor | 58 | 2.60 | 1.76 |
| 6 | B000F6SNPS | Good Earth Herbal Tea, Sweet & Spicy | 143 | 3.69 | 1.76 |
| 7 | B000CRIBCA | High Protein Bars by Think Thin-On the Go | 69 | 3.07 | 1.76 |
| 8 | B000AQJRWG | Tofu Shirataki Noodles Spaghetti Shape | 75 | 3.19 | 1.74 |
| 9 | B002CENRLG | Shirataki Noodles | 101 | 3.54 | 1.72 |
| 10 | B002EDEMLY | Red Vines Red Original Licorice Twists | 77 | 3.69 | 1.73 |

n = number of reviews per product. Mean = mean star rating for each product. SD = standard deviation for product ratings.

The top two factors for polarization were (a) formulation changes in which a product initially liked was changed and got negative reviews and (b) diverse views about the taste of the product. For instance, for formulation change, consumers objected to increases in sugar in a formerly beloved cereal. Illustrative phrases that highlight the opinion diversity were manually extracted and are listed in Table 4.

**Table 4:** Examples of polarized taste comments about the same product

| Abbreviated Product Name | Positive Taste | Negative Taste |
| --- | --- | --- |
| Special K Cereal, Protein | 'taste[s] pretty darn good' | 'it tastes totally horrible!!' |
| blk Premium Alkaline Water | 'it taste[s] just like water with no funny after taste' | 'this stuff tastes like dirty coins' |
| Enjoy Life Chewy Bars | 'the bars have a good taste' | 'tastes like cardboard. zero flavor' |
| Just the Cheese Popped Cheese | 'I think they taste great' | 'worst tasting stuff that I have ever eaten' |
| Good Earth Herbal Tea | 'it tastes fantastic' | 'sick with disappointment over degraded and truly unrecognizable taste' |
| High Protein Bars | 'They taste more like a de[s]sert' | 'tastes of a mixture of cardboard and chemicals' |
| Tofu Shirataki Noodles | 'Tastes identical to real pasta' | 'gross in tast[e] and texture' |
| Shirataki Noodles | 'tasted fresh and enticing each time' | 'taste like rotten fish' |
| Red Vines | 'tasted delicious' | 'they have a horrible after taste' |

## Discussion

The field of ‘Sensory nutrition’ brings together knowledge and methods from sensory, physiology and nutrition sciences to understand the key drivers of nutrient choices so that we can modulate diet to promote human health. Taste is often described as a primary influence on human food selection and intake in nutrition and biopsychology research, e.g., [6]. Studies such as the one just referenced rely on data from several thousand people, but here we demonstrate the primacy of taste among nearly half a million respondents who are unaware of the import of their commentary, essentially catching consumers responding in an unself-conscious way. These results demonstrate that when consumers write about food, rather than cost or its nutritional benefits, they write about taste. Taste is often applied generically to the flavor of food, which has more inputs from different sensory systems, such as the somatosensory system (texture) and the olfactory system (smell) [7]. Therefore, these results could be construed broadly to apply to food flavor, not to taste as narrowly defined by sensory biologists.

### Taste qualities

We learned that *sweet* was the taste quality mentioned most often, almost three times more often than *bitter*, the next closest word used, followed distantly by *sour* and *salty*. This result was a surprise because the opinions about bitterness would be complex and worthy of mention in food product reviews, either as a desirable feature, perhaps in coffee, or an undesirable feature in foods that do not normally taste bitter. However, perhaps bitterness is so rarely present in commercial foods that it is rarely mentioned. Likewise, it was surprising how little saltiness was mentioned by reviewers, given the global attention to salt reduction for health [8]. While the overconsumption of sugar and salt are common public health concerns ([8–10]), consumers have much more to say about sweetness than saltiness, at least in this particular venue. We have learned from analysis of the sweet receptor that different versions arise from inborn genotype, and some people are more sensitive to sweet taste than others because of their genotype [11–15]. With that point in mind, it is interesting that consumers complained more about products being ‘too sweet’ rather than ‘not sweet enough’, indicating that the over-sugaring of processed foods is undesirable for some people and that offering a range of sweetness of products might be even more important that previously realized.

### Polarizing products

We were also interested in polarizing products—those rated variably by different reviewers. While reading these reviews, we noted several trends that appeared to account in part for the polarized ratings. One issue was formulation change—if consumers had bought a product in the past and been satisfied with it, only to find on repurchasing that the formulation had changed (e.g., increasing the product’s sugar content), they down-rated the product. In some ways, these formulation changes muddy the analysis, because the consumers are rating two different products listed with the same product ID.

Often the diversity in viewpoint appeared to arise from different perspectives on a product’s taste. Some reviewers extolled the desirable taste of a particular product whereas others disliked it, sometimes going so far as to berate other reviewers for their opinions. One prominent example was the diversity of viewpoint on product sweetness, which is supported by laboratory-based studies of sweet likers and dislikers, e.g., [16]. Recent studies of personal differences in the liking of intensely sweetened foods suggest this may be an inborn trait [17]

Smell also contributed to the ratings of a few polarizing foods. There are other long-standing debates about the desirable odor of some foods, a common Internet trope being the dislike of cilantro [18,19]. We also noted that smell figured prominently as a polarizing agent for some products with a fishy odor, and we wonder whether the inborn variation in the ability to smell the fishy odor trimethylamine might account for this diversity of viewpoint [20]. Although this study does not allow us to match genotype to reviewer to understand whether these taste disagreements owe to genetics, it does suggest that there is polarization and has identified a handful of products that might be most profitably explored further for genetic effects.

### Limitations

This study has several limitations. It is an analysis of data offered by the online retailer that are freely available to all via a website that encourages exploratory data analysis of large data sets (Kaggle). As such, we had no control over the collection of the data, the number and type of variables included, or the accuracy of the data itself. Thus, all results must be interpreted with this limitation in mind. Further, additional information such as item category or other classifiers would have been useful to limit the analysis to only certain types of foods or to add food type in the analysis. We also learned that foods can have the same identifier but when the manufacturer changes the formulation (e.g., adds sugar) it may lead to polarization because people who preferred the previous version of the product are now dissatisfied. This type of polarization does not arise from diversity of viewpoint about the same food item and these instances dilute the true polarizing response.

A second limitation was the limited choice of words and phrases to count in the reviews, which capture few instances of related speech. While we used a variety of common phrases to capture the concept of sweetness (too much or not enough), many reviewers might have used different words to convey the concept of ‘too sweet’. Also, some phrases about sweetness we classified as ‘neutral’ might be interpreted in one of several ways, e.g., ‘on the sweet side’. Capturing the intended meaning in text strings from real-world situations is imperfect, even using a large palette of terms to describe a certain situation, and those limitations are present here.

Finally, there is an imprecision in the focus on taste, which encompasses several qualities for the average person [7]. The reviewers can use this word both strictly, to evaluate the taste but not smell or texture of the product, or more generally as a holistic quality (for example, ‘…wonderful smell and flavor makes my coffee syrup taste like hazelnuts…’ describes a smell, but not a taste, and is flagged as both in our analysis). This limitation, the imprecision of the word ‘taste’, is offset by the value of capturing real-world perceptions of foods, by people who can report on whatever features they consider to be most important.

### The future

This first study of food reviews from an on-line retailer, taste and polarizing foods portends several avenues of the future research in sensory nutrition. Here we analyzed the content of almost half a million reviews, but it is clearly possible and desirable to perform a similar study on larger cohort, but one in which other information was available about the reviewers, such as demographics, e.g., age and sex, other social and demographic information (e.g., amount of formal education), medical history (e.g., diabetes, hypertension), genotype, and other biological information, e.g., hormone concentrations in the blood or brain imaging. Large scale studies that make the marketplace a laboratory become technically more possible with the potential for linking food purchasing information with electronic medical records. While the social, political and ethical barriers to these types of studies may be insurmountable, studies at this scale are increasingly technically possible. These studies could reveal previously overlooked patterns of food consumption and disease and point to new avenues of biology to explain why people choose the foods they do.

### Conclusion

We learned from these data that, when it comes to commercially available food products, taste matters and that there are diverse viewpoints about some products that may stem in part from differences in basic biology. Looking ahead and drawing on the research steps from the preceding paragraph, it may be possible to find genotypes (for instance, in taste receptors) that predict who will or will not like the taste of a given product. This idea may translate into healthier foods, if producers can reduce sugar or salt in ways that appeal to groups that prefer those products, by linking genetics (taste-related genotypes) to behavior (food-purchasing habits). This study is a step toward realizing this idea.

## Electronic resources

https://www.kaggle.com/snap/amazon-fine-food-reviews

https://github.com/joelmainland/Taste_is_king

## Acknowledgments

This study was funded by institutional funds from the Monell Chemical Senses Center. We thank members of R-Club who provided coding and analysis advice: Cailu Lin, Stephanie Gervasi, Molly Spencer, Steven Brooks, Alissa Nolden, Marissa Karmarck, Nicolle Murphy, Carolyn Novaleski, Genevieve Bell, and Emily Mayhew. Nancy Rawson commented on a draft of the manuscript. We thank two anonymous reviewers for their insightful comments.

---

**Supplemental Table 1.**
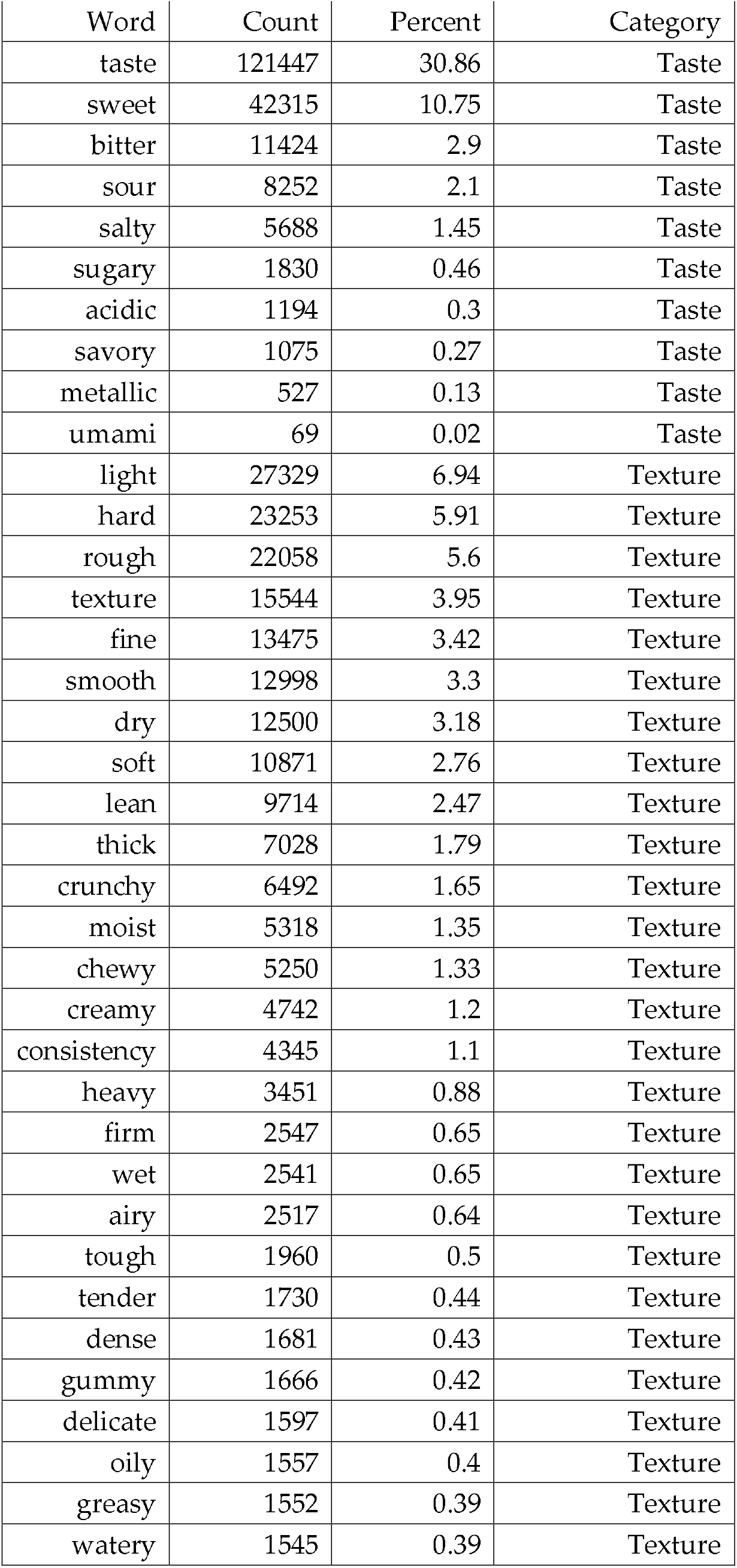
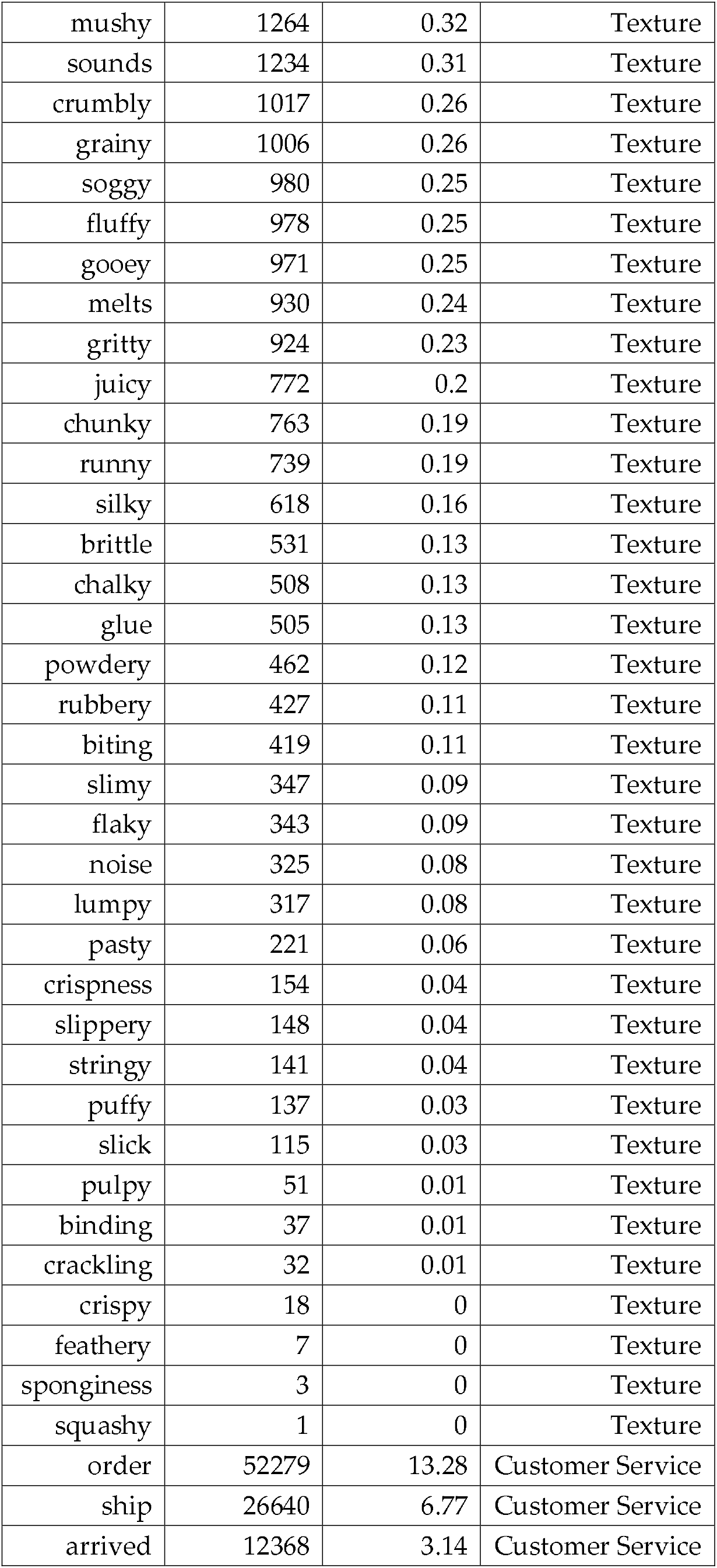
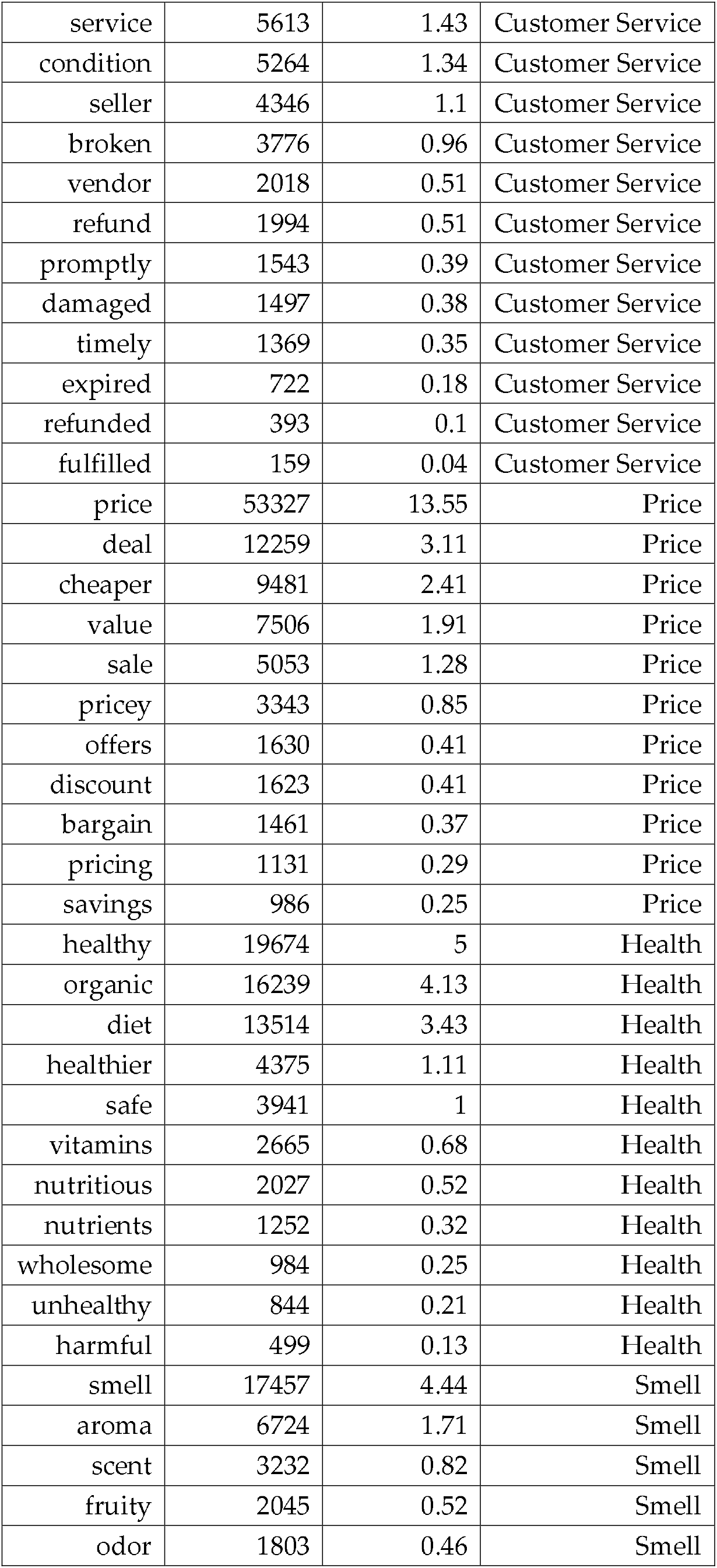
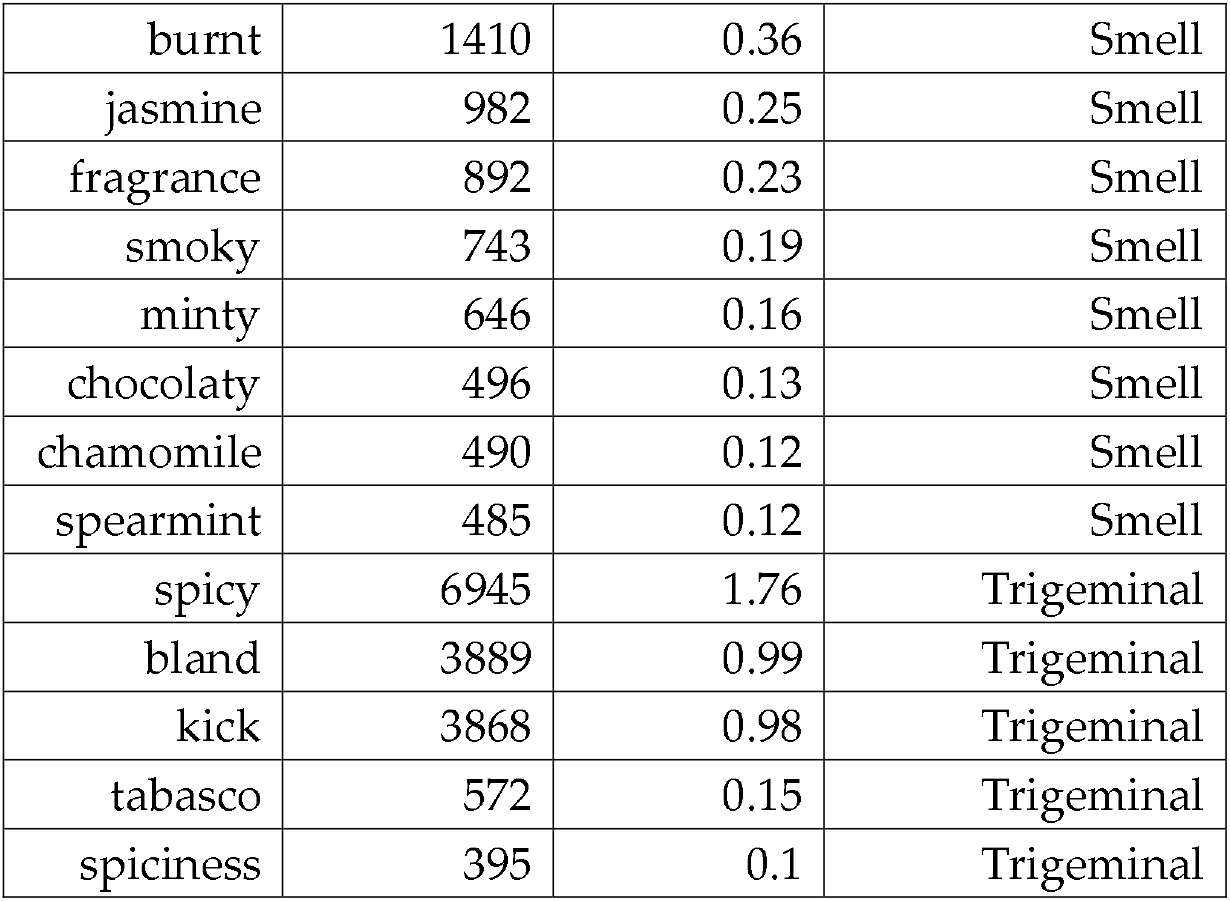
Word list from Word2Vec analysis

**Supplemental Table 2:** Sweet phrases

| Phrase | Count | Percent | Type |
| --- | --- | --- | --- |
| not very sweet | 163 | 0.04 | Under |
| not sweet enough | 94 | 0.02 | Under |
| not sweet enough for me | 11 | 0 | Under |
| too sweet | 3121 | 0.79 | Over |
| very sweet | 1297 | 0.33 | Over |
| to sweet | 1020 | 0.26 | Over |
| overly sweet | 616 | 0.16 | Over |
| so sweet | 378 | 0.1 | Over |
| really sweet | 230 | 0.06 | Over |
| sickly sweet | 141 | 0.04 | Over |
| sickeningly sweet | 114 | 0.03 | Over |
| cloyingly sweet | 108 | 0.03 | Over |
| pretty sweet | 106 | 0.03 | Over |
| overwhelmingly sweet | 38 | 0.01 | Over |
| excessively sweet | 21 | 0.01 | Over |
| overpoweringly sweet | 20 | 0.01 | Over |
| disgustingly sweet | 13 | 0 | Over |
| ultra sweet | 6 | 0 | Over |
| couldn't stand how sweet it was | 1 | 0 | Over |
| not too sweet | 2550 | 0.65 | Neutral |
| slightly sweet | 855 | 0.22 | Neutral |
| not overly sweet | 700 | 0.18 | Neutral |
| sweet enough | 644 | 0.16 | Neutral |
| on the sweet side | 212 | 0.05 | Neutral |
| not to sweet | 135 | 0.03 | Neutral |
| not so sweet | 83 | 0.02 | Neutral |
| sweet enough for me | 36 | 0.01 | Neutral |
| not overwhelmingly sweet | 28 | 0.01 | Neutral |
| not cloyingly sweet | 27 | 0.01 | Neutral |
| not sickly sweet | 26 | 0.01 | Neutral |
| not sickeningly sweet | 22 | 0.01 | Neutral |
| not overpoweringly sweet | 18 | 0 | Neutral |
| not really sweet | 15 | 0 | Neutral |
| not excessively sweet | 10 | 0 | Neutral |
| not disgustingly sweet | 4 | 0 | Neutral |
| sweet as i would prefer | 2 | 0 | Neutral |
| not ultra sweet | 2 | 0 | Neutral |
| not on the sweet side | 1 | 0 | Neutral |

The count refers to the number of reviews that contain the relevant phrase and the percent of all reviews that contain that phrase. Type refers to undersweetness, oversweetness or neutral comments.

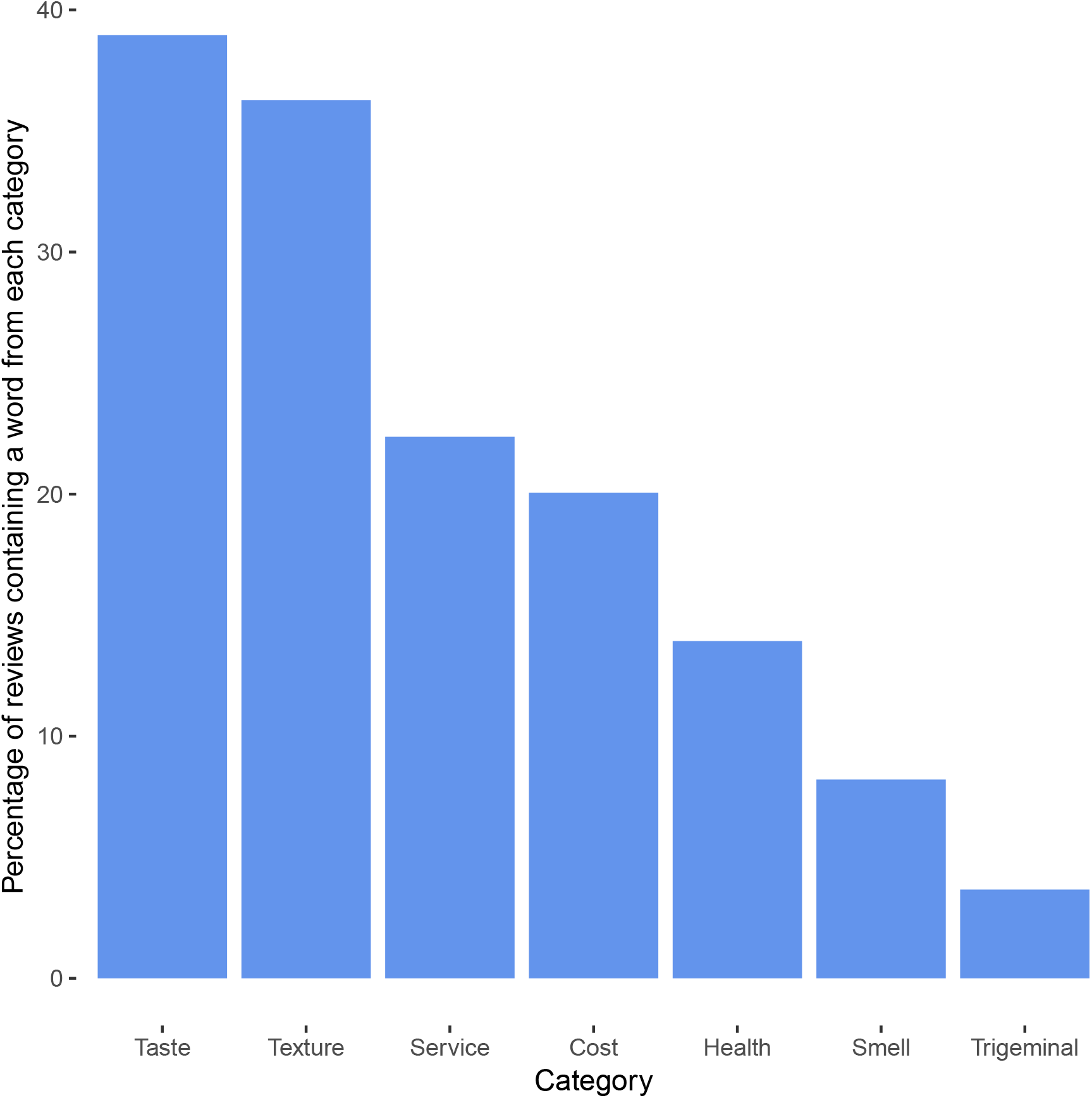

